# Immobilization-free chemotaxis analysis reveals the novel behavioral mode of “leaving” in *Caenorhabditis elegans*

**DOI:** 10.64898/2026.07.01.734387

**Authors:** Shiori Onoue, Koji Kyoda, Shuichi Onami

**Affiliations:** Graduate School of Frontier Bioscience, The University of Osaka, Suita, Osaka 565-0871, Japan; Laboratory for Developmental Dynamics, RIKEN Center for Biosystems Dynamics Research, 2-2-3 Minatojima-minamimachi, Chuo-ku, Kobe, Hyogo 650-0047, Japan

## Abstract

Animals balance staying in a favorable environment with exploring new ones. In *C. elegans* chemotaxis, the process by which worms migrate toward an attractant has been extensively studied. However, what happens after they reach it remains largely unexplored, partly because conventional assays immobilize worms at the point of arrival. Here, we quantitatively analyzed chemotactic behavior upon reaching an attractive odor source using an immobilization-free chemotaxis assay. We observed that 62% animals left the isoamyl alcohol region after initially approaching it, a behavior we termed “leaving behavior.” Quantitative analysis revealed that leaving behavior represents a distinct locomotor state compared with free-moving, high-concentration odor avoidance, and approach behavior. To test whether leaving behavior is related to olfactory adaptation, we analyzed mutants in adaptation-related genes. The proportion of leaving behavior was significantly increased in *egl-4* loss-of-function mutants compared with wild-type animals, whereas arr-1 mutants showed no significant difference. These results suggest that *egl-4* negatively regulates leaving behavior, suggesting a role for this kinase in stabilizing post-arrival behavioral states beyond its known function in olfactory adaptation. Our findings indicate that chemotaxis involves dynamic behavioral transitions even after reaching an attractant, consistent with an exploration–exploitation trade-off framework.

**Significance Statement:** By eliminating immobilization from a conventional chemotaxis assay, this study reveals behaviors that are typically obscured after animals reach an attractant. In *C. elegans*, we identify a post-arrival behavioral mode, “leaving,” in which animals move away from an attractive odorant (isoamyl alcohol) after initially approaching it. Leaving behavior is quantitatively distinct from free-moving behavior, high-concentration odor avoidance, and approaching behavior. We further show that the frequency of leaving behavior is increased in animals lacking the *egl-4* gene. These findings extend chemotaxis analysis beyond the point of odor source arrival and suggest that the nervous system actively drives behavioral switching even after a goal is reached, broadening our understanding of how sensory circuits govern exploration–exploitation decisions.

## Introduction

Animals must balance staying in a favorable environment with exploring new ones. Many animals navigate their environment by processing odor information (Adler, 1969; Brenner, 1974; Brown and Berg, 1974; Berg, 1975; Porter et al., 2007; Baker et al., 2018). In *Caenorhabditis elegans*, chemotaxis has been extensively studied with respect to which odorants are detected, which sensory neurons and receptors mediate their detection, and whether odor presentation elicits attraction, avoidance, or no response (Ward, 1973; Bargmann and Horvitz, 1991; Bargmann et al., 1993; Pierce-Shimomura et al., 2004; Bargmann, 2006; Yoshida et al., 2012; Tanimoto et al., 2017). Behaviors such as approaching an attractant and avoiding a repellent have also been characterized in detail (Iino et al., 2009). However, what *C. elegans* individuals do after they reach an attractant remains much less clear.

Early studies noted that after approaching an attractant *C. elegans* individuals remain near the source for a period and then leave, sometimes repeatedly leaving and reapproaching. This behavior was proposed to reflect habituation or adaptation to the odor (Ward, 1973; Bargmann and Horvitz, 1991).

Because these behaviors occur after *C. elegans* individuals reach the attractant, they are difficult to analyze in conventional chemotaxis assays, which commonly use sodium azide to immobilize the nematodes at the odor source. This assay design has important advantages, including high quantification and experimental efficiency, such that it has been widely adopted (Bargmann et al., 1993; Margie et al., 2013; Queirós et al., 2021). In contrast, experimental designs that use anesthetics make it difficult to continuously observe behavior after arrival at an attractant. As a result, analyses of chemotactic behavior have primarily focused on the process of reaching the odor source, whereas post-arrival behaviors may not have been examined in sufficient detail.

*Caenorhabditis elegans* adapts to attractants following prolonged exposure (Thompson and Spencer, 1966; Colbert and Bargmann, 1995). Among the genes implicated in olfactory adaptation, *egl-4* and *arr-1* play important roles. In pre-exposure assays, wild-type individuals typically show reduced chemotaxis after odor exposure, whereas *egl-4* and *arr-1* loss-of-function mutants retain chemotaxis (Colbert and Bargmann, 1995; Lee et al., 2010; Merritt et al., 2022). However, such adaptation assay designs do not reflect the conditions *C. elegans* individuals encounter in natural situations. In natural environments, individuals experience an attractive odor continuously and decide whether to remain or leave based on that ongoing sensory input.

Thus, behaviors that occur after reaching an attractant have remained largely unexplored, in part because of experimental constraints. Here, we analyzed chemotactic behavior in the absence of sodium azide, thus allowing continuous observation of *C. elegans* behavior after arrival at the odor source, while also using computational and genetic approaches to investigate post-arrival behavior.

## Results

### Leaving behavior occurs after approaching the attractant

Previous studies have described nematodes leaving the attractant after approaching it. To confirm this phenomenon, we performed a conventional population chemotaxis assay using isoamyl alcohol as the odor source (Fig. 1A). To observe *C. elegans* individuals’ behavior after reaching the attractant, we omitted an anesthetic and captured five snapshots at 10-min intervals (Fig. 1B).

**Figure 1.**
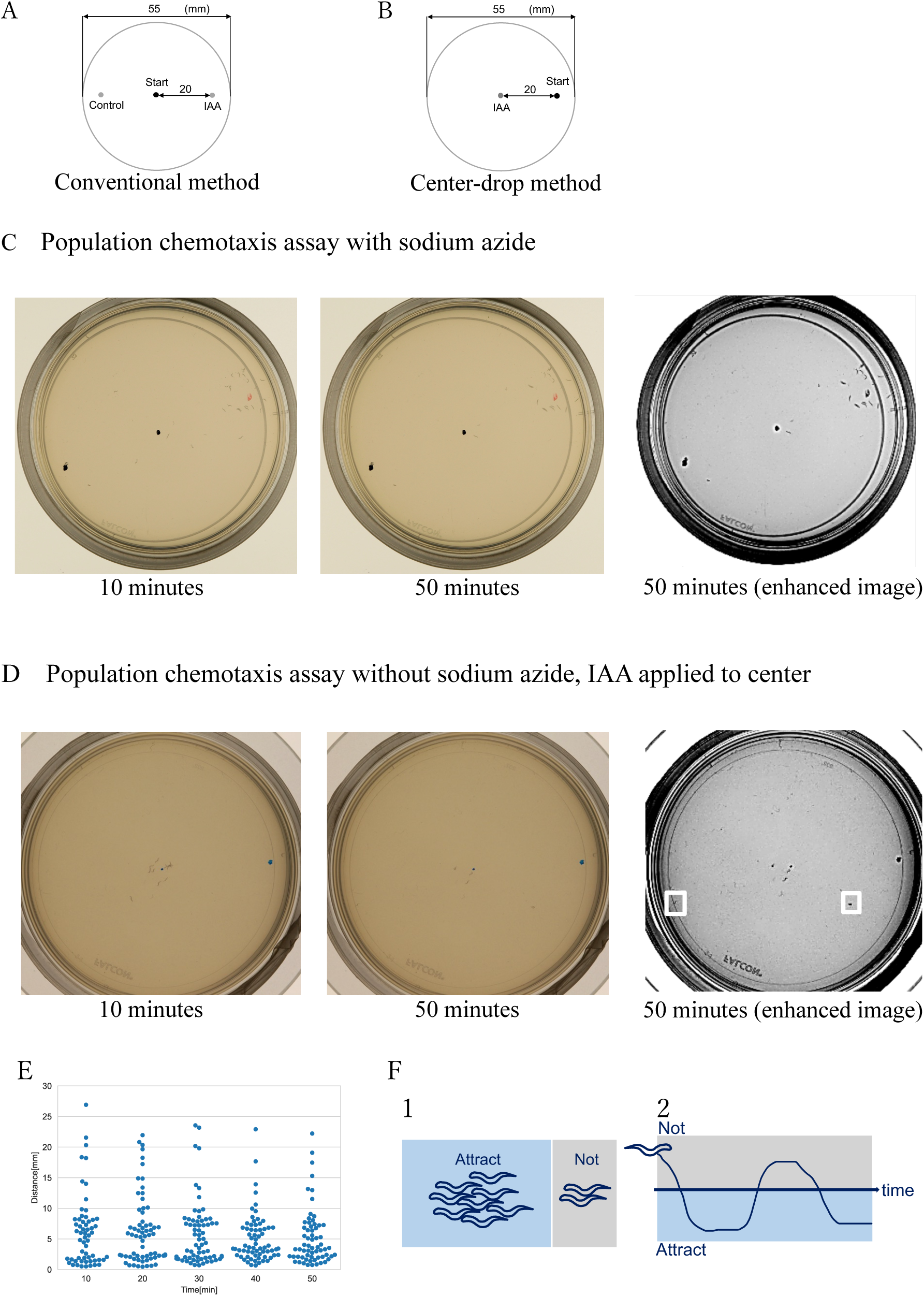
Comparison of conventional and immobilization-free chemotaxis assays (A) Conventional chemotaxis assay method with isoamyl alcohol (IAA) as the attractant. (B) Center-drop method: An attractant was placed at the center of the plate without sodium azide. (C) Representative images of a conventional chemotaxis assay with sodium azide at 10 and 50 min after assay initiation. (D) Representative images of the immobilization-free chemotaxis assay. White rectangles indicate individuals located far from the center of the plate. (E) Distance between each individual and the IAA drop point in the population chemotaxis assay (N = 48). (F) Two possible explanations for individuals located far from the IAA source: (1) Individuals were not attracted to IAA or (2) individuals were in a behavioral state that prevented an approach to IAA.

Although many individuals aggregated near the odor source, a subset of individuals tended to be located away from it (Figs. 1A, C). We confirmed a similar tendency when the population assay was performed using a central-drop method in which isoamyl alcohol was placed at the center of the plate (Figs. 1D, E; Fig. S1E).

To determine whether the population included both attracted and unattracted individuals and/or whether the same individuals repeatedly approached and left the attractant (Fig. 1F), we manually quantified locomotor trajectories. We observed that all *C. elegans* (*n* = 8) approached the odor source, and 3 individuals subsequently left the isoamyl alcohol source during the 15-min observation (Fig. 2A). Leaving after attraction was also observed in the individual chemotaxis assay (Fig. 2B; Movie S1). Based on these results, we confirmed that some nematodes repeatedly approach isoamyl alcohol and exhibit leaving behavior. Because nematodes can be influenced physically and physiologically by other individuals, we used the individual chemotaxis assay for subsequent analyses to exclude such effects.

**Figure 2.**
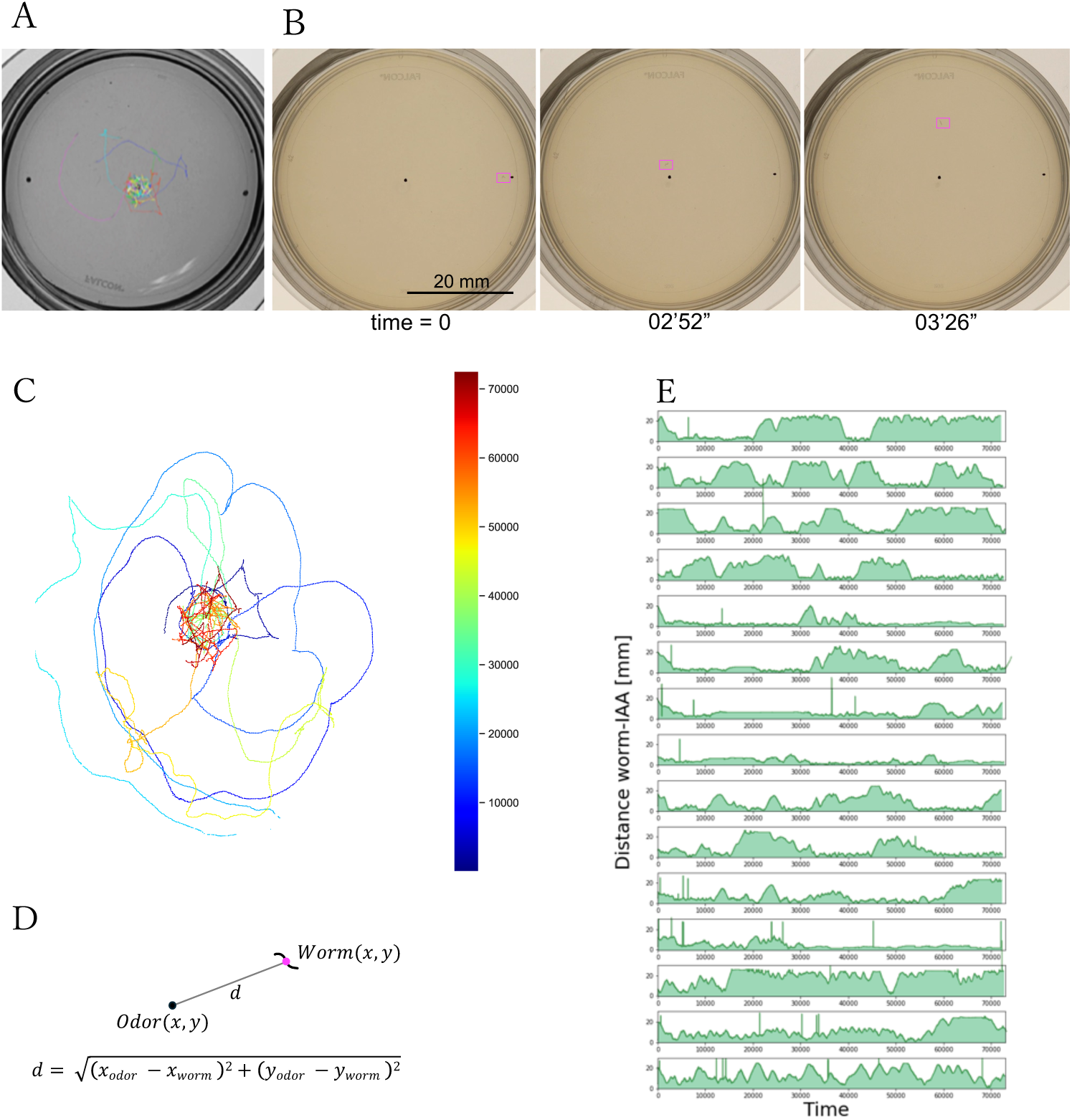
Behavioral trajectories of individual nematodes after approaching an attractant (A) Population chemotaxis behavior was recorded, and behavioral trajectories were tracked manually. Each color represents the behavioral trajectory of a single individual. Chemotaxis started from the black dot at the right. Single-nematode chemotaxis behavior was examined using the original chemotaxis method. A movie was recorded for 50 min. Chemotaxis started from the black dot at the right, and magenta rectangles indicate the location of the worm (see Supplemental video 1). (C) The trajectory of chemotaxis behavior, with a 50-min single chemotaxis assay using the center-drop method. Dark blue indicates the start time point of chemotaxis, and red indicates the end of the assay. (D) Equation for Euclidean distance d between the nematode location and the isoamyl alcohol drop point. (E) Under isoamyl alcohol conditions, for each individual distance d was measured and plotted against time.

To statistically analyze behavior on the plate, we developed Simple Worm Tracker (SWT) (Fig. S2), software that measures locomotor trajectories using background subtraction (Piccardi, 2004; Zivkovic, 2004). Using SWT, we analyzed 50-min chemotaxis videos to quantify approaching and leaving (Fig. 2C; Fig. 1B). From the resulting trajectory data, we measured fluctuations in the distance (*d*) from each *C. elegans* individual to the isoamyl alcohol source (Fig. 2D). We found that the leave phenomenon occurred in 62% (31/50) of animals after they approached the isoamyl alcohol. The timing of switching from approach to leave, as well as the frequency of switching, varied widely among individuals (Fig. 2E). We refer to nematodes leaving the odor source after approaching isoamyl alcohol as “Leaving behavior.”

### Leaving behavior is distinct from free-moving behavior

A simple explanation for the observed phenomenon is that, after exhibiting positive chemotaxis toward the isoamyl alcohol source, the nematode’s attraction dissipates and it returns to unstimulated, free-moving, resulting in an increased distance from the odor source.

To examine whether locomotion during leaving behavior is equivalent to locomotion during free-moving behavior, we quantitatively compared the locomotion traits of *C. elegans* individuals during leaving behavior with those during free-moving behavior. Visual inspection suggested that long, sustained runs were more characteristic during leaving behavior than during free-moving behavior. We therefore compared sharp-turn frequency, average speed, and trajectory length between leaving behavior and free-moving behavior. Leaving behavior was defined as a run that originated within 5 mm of the isoamyl alcohol source and continued until the nematode reached a position at least 7 mm away from the source (Fig. 3A; Figs. S3A–C; Figs. S4A–C; see Methods).

**Figure 3.**
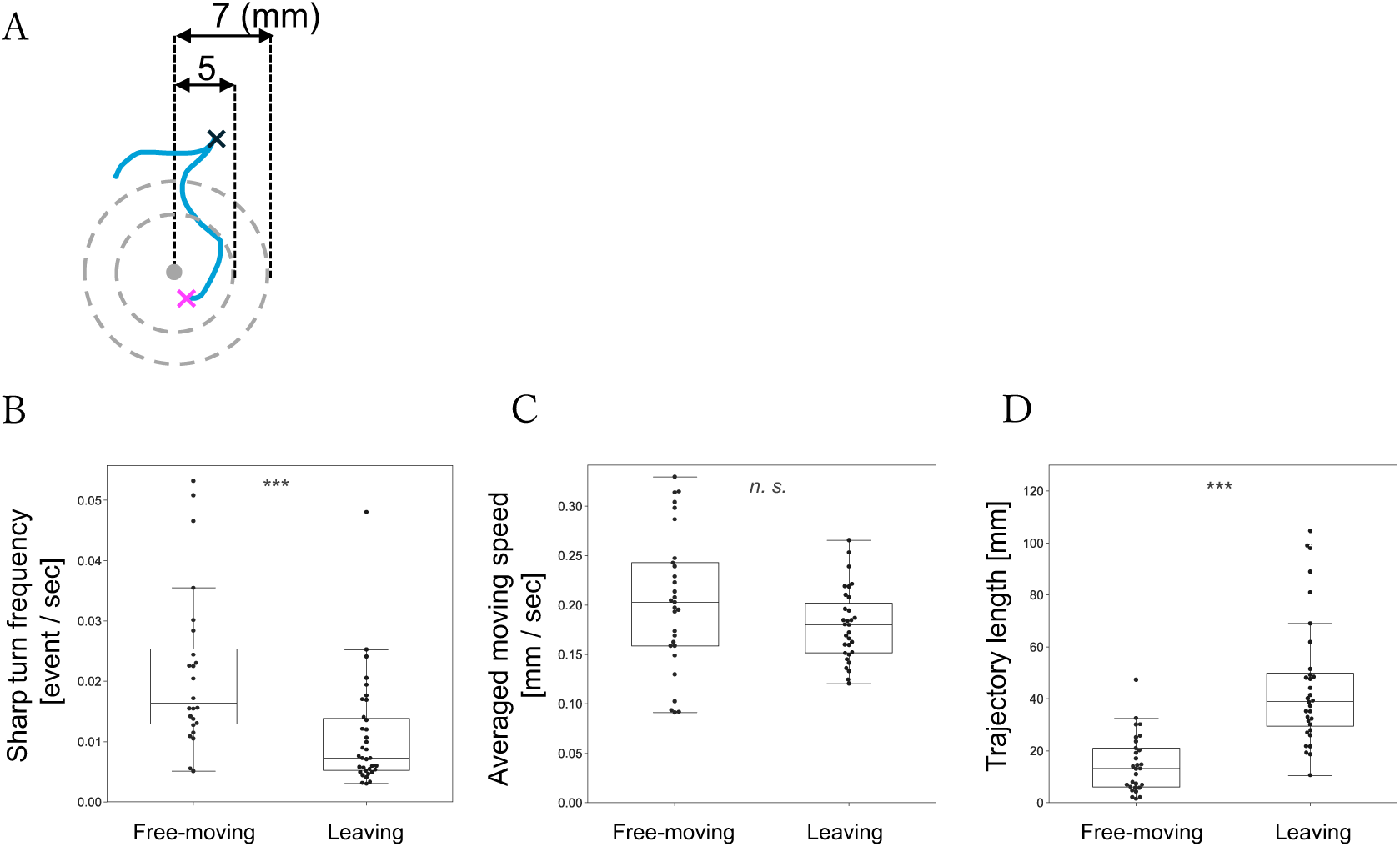
Definition and locomotor features of leaving behavior compared with free-moving behavior (A) Definition of the leaving behavior. The gray dot represents the isoamyl alcohol drop point. The magenta cross represents the initial point, and the black cross represents the endpoint of the leaving behavior (Figure S3, S4, method). The same criteria were used to define free-moving behavior during non-stimulated 50-min periods. Behavioral features were compared between the free-movement (n = 24) and the leaving behavior (n = 35): (B) sharp-turn frequency, (C) average speed, (D) trajectory length. Significant differences based on the Mann-Whitney U-test are noted as *p < 0.05, **p < 0.01, ***p < 0.001.

We found that, compared with free-moving behavior (Fig. 3A), leaving behavior exhibited a significantly lower sharp-turn frequency (Mann–Whitney U test, *p* = 0.000152, Fig. 3B) and a significantly longer trajectory length (Mann–Whitney U test, *p* = 0.0000000281, Fig. 3D), whereas mean locomotor speed did not differ significantly (Mann–Whitney U test, *p* = 0.12037, Fig. 3C). Based on these results, we concluded that leaving behavior is distinct from free-moving behavior.

### Leaving behavior is distinct from high-concentration isoamyl alcohol avoidance behavior

Avoidance behavior in response to high-concentration isoamyl alcohol is a well-documented example of odor-evoked behavior in which nematodes move away from a chemical stimulus. To examine whether locomotion during leaving behavior is equivalent to locomotion during high-concentration isoamyl alcohol avoidance behavior, we quantitatively compared these behaviors.

We found that the leaving behavior exhibited a significantly lower sharp-turn frequency and a significantly longer trajectory length than high-concentration isoamyl alcohol avoidance behavior (Mann–Whitney U test, *p* = 0.00379, Fig. 4B; Mann–Whitney U test, *p* = 0.000116, Fig. 4D). Average speed did not differ significantly (Mann–Whitney U test, *p* = 0.16662, Fig. 4C). Based on these results, we concluded that leaving behavior is distinct from high-concentration isoamyl alcohol avoidance behavior.

**Figure 4.**
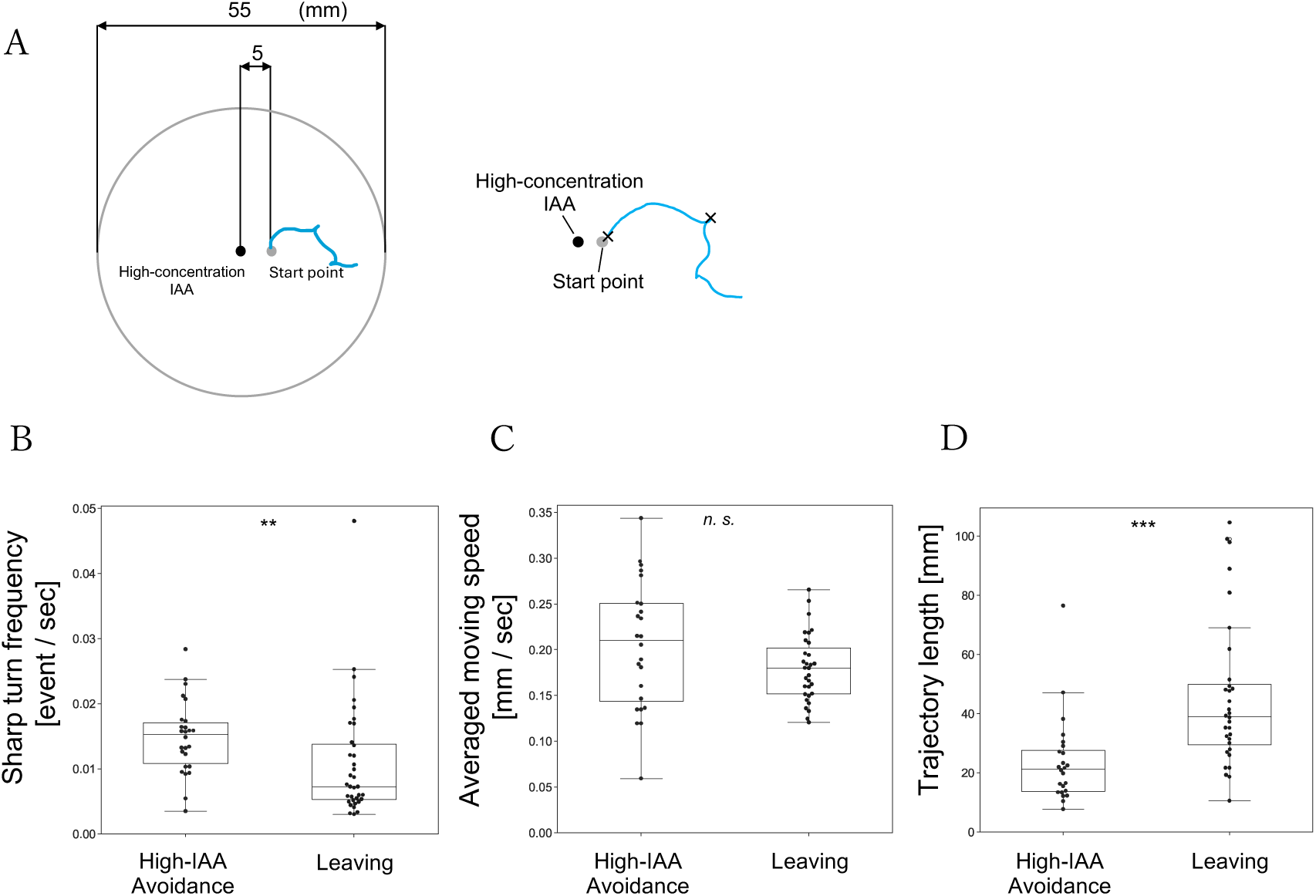
Locomotor features of leaving behavior compared with high-concentration isoamyl alcohol avoidance behavior (A) In the high-concentration isoamyl alcohol avoidance behavior assay, 2 μL of undiluted isoamyl alcohol was placed at the center of an NGM plate, and the position 5 mm away from the drop point was designated as the nematode behavior start point. After washing, a single nematode was selected from the solution (see Method), placed at the designated start point, and immediately filmed for 10 min. Behavioral trajectories were quantified using our developed software, Simple Worm Tracker. The trajectory from the behavior start point to a subsequent sharp turn was extracted as high-concentration isoamyl alcohol avoidance behavior (n = 26) and leaving behavior (n = 35). (B) Sharp-turn frequency, (C) average speed, (D) trajectory length. Significant differences based on the Mann-Whitney U-test are noted as *p < 0.05, **p < 0.01, ***p < 0.001.

### Leaving behavior is distinct from approaching behavior

Another well-characterized odor-guided behavior is approaching behavior toward isoamyl alcohol (e.g., Bargmann et al., 1993). To examine whether locomotion during leaving behavior is equivalent to locomotion during approaching behavior, we quantitatively compared these behaviors.

We found that the leaving behavior exhibited a significantly lower sharp-turn frequency, a significantly slower average speed, and a longer trajectory length than approaching behavior (Mann–Whitney U test, *p* = 0.00000365, Fig. 5A; Mann–Whitney U test, *p* = 0.01579, Fig. 5B; Mann–Whitney U test, *p* = 0.000000690, Fig. 5C). Based on these results, we concluded that leaving behavior is distinct from approaching behavior.

**Figure 5.**
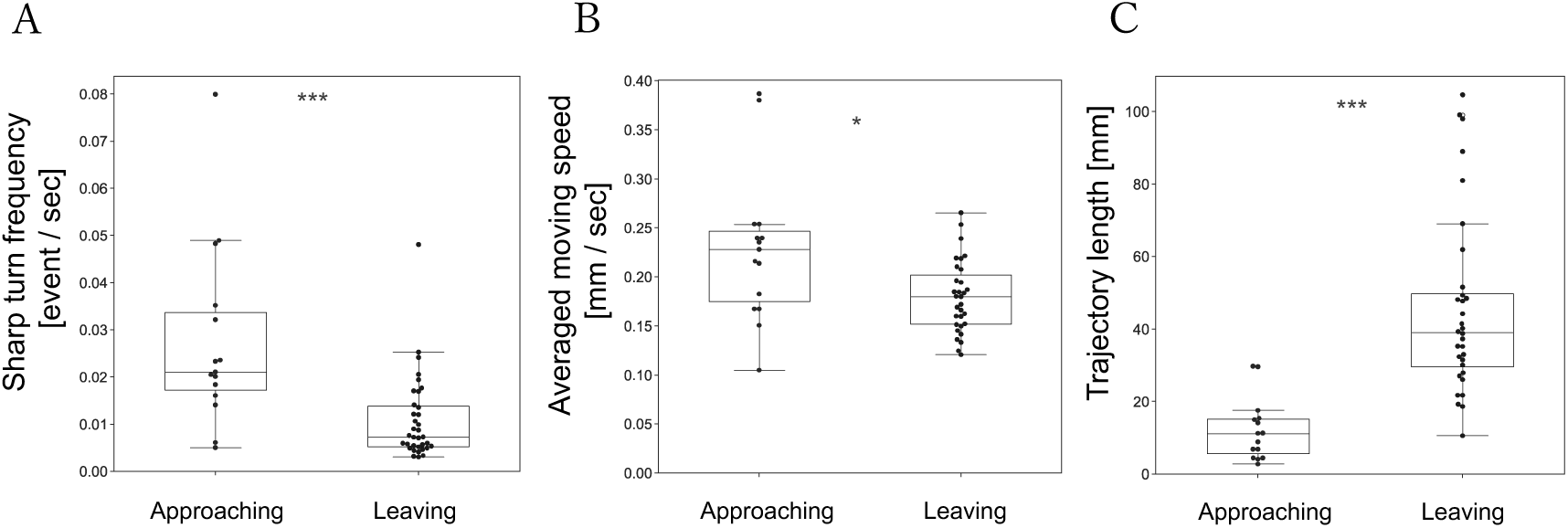
Locomotor features of leaving behavior compared with approaching behavior Behavioral features were compared between approaching (n = 15) and leaving behavior (n = 35): (A) sharp-turn frequency, (B) average speed, (C) trajectory length. Significant differences based on the Mann-Whitney U-test are noted as *p < 0.05, **p < 0.01, ***p < 0.001.

### Leaving behavior is not an avoidance response to low-concentration isoamyl alcohol

Does low-concentration isoamyl alcohol trigger an avoidance-like response under sudden stimulation conditions? To examine whether leaving behavior represents an avoidance response to low concentrations of isoamyl alcohol, we analyzed nematode behavior while maintaining the experimental setup used for high-concentration isoamyl alcohol avoidance behavior, only changing the concentration of isoamyl alcohol to 1/100.

We found that under the 1/100 isoamyl alcohol condition, all *C. elegans* individuals approached the odor source to within 5 mm, whereas under the undiluted isoamyl alcohol condition, no individuals approached the odor source (Figs. 6A–C). After approaching the 1/100 isoamyl alcohol source, 5 of 15 individuals exhibited leaving behavior (Fig. 6A). Based on these results, we concluded that leaving behavior is not an avoidance response to low concentrations of isoamyl alcohol.

**Figure 6.**
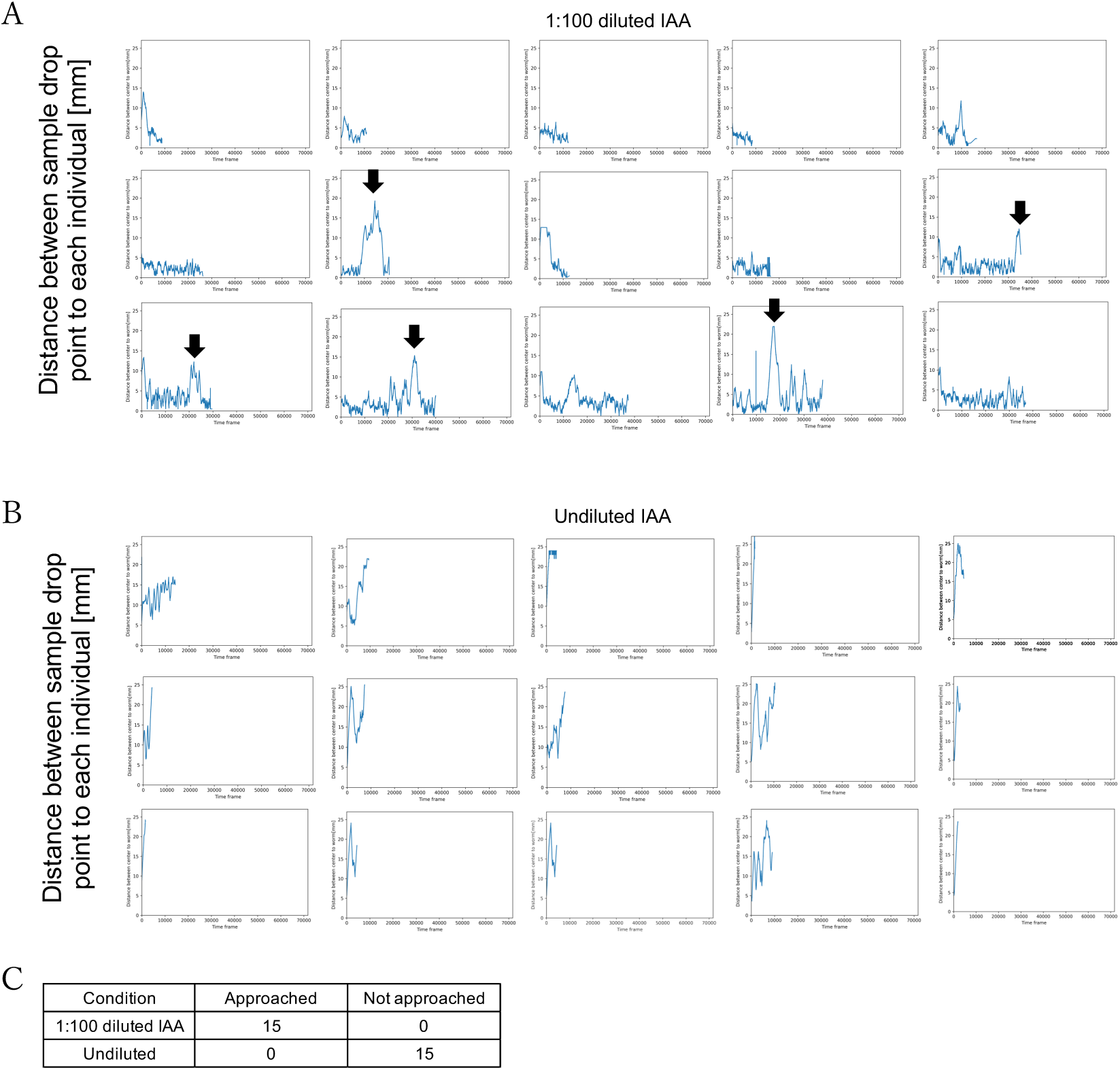
Extent of approach, leaving, and avoidance behaviors in response to low and high concentrations of isoamyl alcohol (A) Temporal variation in distance from the 1:100 isoamyl alcohol (IAA) drop point to each individual. Black arrows indicate actions that appear to be leaving behavior. (B) Temporal variation in the distance from the undiluted IAA drop point to each individual. (C) For each condition, the number of individuals that approached within 5 mm of the IAA drop point and the number that did not approach.

### egl-4 suppresses leaving behavior

We hypothesized that adaptation to isoamyl alcohol following prolonged exposure causes leaving behavior. To test this hypothesis, we compared the proportion of leaving behavior among *egl-4* and *arr-1* loss-of-function mutants and wild-type individuals. The genes *egl-4* and *arr-1* promote adaptation to odor stimuli in *C. elegans* (Pierce and Lefkowitz, 2001; Fujiwara et al., 2002; L’Etoile et al., 2002; Hino et al., 2021; Merritt et al., 2022).

Nematodes were subjected to a 50-min chemotaxis assay. Their responses were classified as “approached” if they reached the isoamyl alcohol source, “leaving” if they exhibited leaving behavior at least once after approaching, or “staying” if they remained near the odor source. The proportions of individuals in each category were compared across genotypes. We found that in *egl-4* loss-of-function mutants (MT1073), the proportion of leaving individuals was 93% (29/31), which was significantly greater than that observed in wild-type N2 (62%, 31/50; χ²(1) = 8.34, *p* = 0.00387; Figs. 7A, C). In contrast, the proportion of leaving individuals in *arr-1* loss-of-function mutants (RB660) was 75% (21/28) and did not differ significantly from that of N2 (χ²(1) = 0.84, *p* = 0.35863; Figs. 7A, C). Loss of adaptation-related function did not lead to a reduction in the occurrence of leaving behavior. Based on these results, we concluded that *egl-4* plays a suppressive role in regulating leaving behavior.

**Figure 7.**
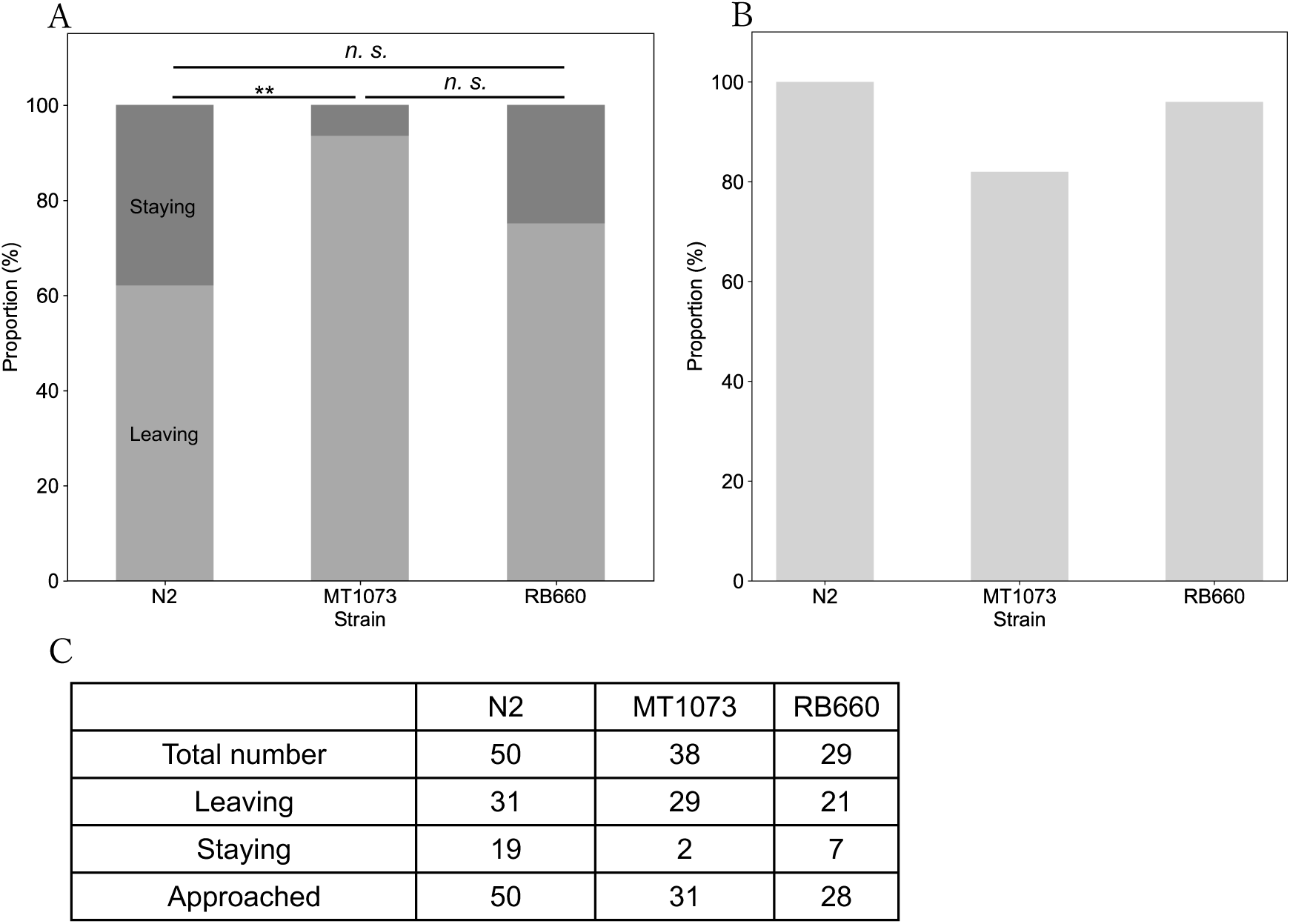
Proportion of individuals exhibiting leaving behavior in wild-type and adaptation-related gene mutants (A) Among the individuals that approached isoamyl alcohol, those that remained at the site were counted as staying (dark gray) and those that left as leaving (gray); the percentages are shown in bars. (B) Percentage of individuals that approached within 5 mm of the isoamyl alcohol drop point (light gray) among those assayed in chemotaxis experiments. (C) Number of leaving individuals, staying individuals, and approaching individuals in wild-type (N2), egl-4 loss-of-function mutants (MT1073), and arr-1 loss-of-function mutants (RB660). Significant differences based on a chi-square test are noted as *p < 0.05, **p < 0.01, ***p < 0.001.

## Discussion

We identified a novel behavioral mode, termed leaving behavior, that emerges after animals approach an attractant. We further showed that *egl-4* likely contributes to leaving behavior in a suppressive, rather than promotive, manner.

### Leaving behavior represents a behavioral mode distinct from previously described behaviors

We found that 62% of nematodes exhibited leaving behavior after approaching isoamyl alcohol (Fig. 2E). Leaving behavior was characterized by a significantly lower sharp-turn frequency and a longer trajectory length compared with those of free-moving behavior, approaching behavior, and high-concentration isoamyl alcohol avoidance behavior (Figs. 3–5). The most straightforward interpretation of these results is that leaving behavior constitutes an independent behavioral mode that is distinct from other known behaviors in *C. elegans*.

One possible explanation is that leaving behavior reflects avoidance behavior in response to low concentrations of isoamyl alcohol. However, under low-concentration conditions, nematodes did not exhibit the immediate movement away from the odor source that is characteristic of high-concentration isoamyl alcohol avoidance behavior. Instead, all individuals approached the odor source immediately after placement on the plate (Fig. 5A–C). Thus, leaving behavior is unlikely to represent avoidance behavior toward low concentrations of isoamyl alcohol.

### egl-4 suppressively regulates leaving behavior

We initially expected that leaving behavior would be suppressed in the *egl-4* loss-of-function mutant, as *egl-4* is one of the representative genes known to promote adaptation. Contrary to this expectation, we found that the proportion of individuals exhibiting leaving behavior was significantly higher in the *egl-4* loss-of-function mutant than in wild-type individuals, suggesting that *egl-4* plays an inhibitory role in the regulation of leaving behavior.

Previous studies primarily analyzed behavioral outputs in terms of whether *C. elegans* individuals approach an attractant following prior exposure (Colbert and Bargmann, 1995; L’Etoile et al., 2002; Lee et al., 2010; Hino et al., 2021). In contrast, our analysis focused specifically on individuals who had already approached the attractant and examined their subsequent behavioral choice: whether to remain near the attractant or leave. Because the experimental conditions and behavioral context differ, our results do not necessarily refute the hypothesis that adaptation to attractants induces leaving behavior. Rather, our findings suggest that *egl-4* may contribute to adaptation not only by suppressing sensory responses to odor stimuli but also by stabilizing behavior after the nematode reaches an attractant.

The increase in leaving behavior observed in the *egl-4* loss-of-function mutant (Figs. 7A–C) can be interpreted in a manner consistent with the known role of *egl-4* in behavioral regulation. Previous studies reported that *egl-4* loss-of-function mutants are biased toward roaming rather than dwelling (Fujiwara et al., 2002). In *C. elegans*, dwelling is characterized by lower average speed and high turn frequency within a localized food patch, whereas roaming involves higher average speed and lower turn frequency over a broader area (Ben Arous et al., 2009; Flavell et al., 2020). *egl-4* is activated by sensory neuronal inputs from cilia and promotes the dwelling state (Fujiwara et al., 2002). Based on these findings, *egl-4*-loss-of-function mutants are likely biased toward roaming over dwelling, resulting in an internal state in which local staying behavior is unstable. Accordingly, in wild-type individuals, *egl-4* can be interpreted as suppressing leaving behavior by stabilizing staying behavior after the nematode approaches an attractant.

### Adaptation to attractive stimuli and change in the dynamics of behavior

Consistent with previous studies, we initially assumed that leaving behavior represented a free-movement-like locomotor pattern that emerges because of sensory adaptation to an attractive stimulus, in which prolonged exposure reduces the sensitivity of sensory neurons and weakens chemotactic behavior (Colbert and Bargmann, 1995; L’Etoile et al., 2002; Lee et al., 2010). However, quantitative analysis revealed that leaving behavior constitutes a novel behavioral mode distinct from free-moving. Furthermore, analyses using loss-of-function mutants of genes known to promote adaptation suggested that the trigger for leaving behavior involves mechanisms different from those underlying conventional adaptation. The leaving behavior observed in this study seems to represent a change in the dynamics of behavior that occurs while the nematodes continue to sense the attractive stimulus.

Behavioral switching in response to attractive stimuli has been studied within the framework of exploratory behavior, such as switching between staying in and leaving food patches (Hills., 2015). In *Drosophila*, behavioral switches between staying and leaving are known to depend on internal states in environments containing multiple food patches (Corrales-Carvajal et al., 2016). In primates, decisions about whether to continue exploiting the current patch or disengage and switch toward exploration have also been extensively investigated (Hayden et al., 2011; Kolling et al., 2012). The change in the dynamics of behavior after arriving at an attractant identified in this study suggests that chemotactic behavior in *C. elegans*, as well as other forms of animal behavior, should be understood as a dynamic phenomenon involving state transitions rather than as a static behavioral output.

### Advantages and limitations of the Simple Worm Tracker

We developed and employed the behavioral quantification system SWT to continuously analyze nematode behavior during chemotaxis with high temporal and spatial resolution. SWT offers several advantages. First, by continuously tracking nematode centroids at a high frame rate, the software enables the extraction of detailed behavioral trajectories, including spatial positions on the plate. Second, SWT can be applied to high-resolution, high-frame-rate videos, enabling the simultaneous acquisition of behavioral information across multiple spatial scales, from coarse positional data to fine-scale body postures, such as body bending (Fig. S2C). Third, SWT is not restricted to specific experimental conditions and can be applied to diverse paradigms, including approaching behavior, avoidance behavior, and free-movement behavior, making it suitable for comparative analysis of behavioral modes (Swierczek et al., 2011; Panadeiro et al., 2021).

However, SWT also has limitations. It does not support simultaneous tracking of multiple individuals and is therefore optimized for single-nematode analysis. Due to the nature of background-subtraction-based detection, the accuracy of centroid detection decreases when individuals remain stationary. In addition, sharp turns currently must be identified manually, and observer-dependent bias cannot be completely ruled out. In this study, however, sharp turns were defined as events in which an omega turn followed a reversal, as indicated by clear postural changes, making subjective judgment unlikely to vary substantially between observers. Future improvements, such as automated sharp-turn detection using machine learning and extension to multi-worm tracking, are expected to enable more precise analysis of behavioral transitions.

### Future research on leaving behavior

By continuously monitoring chemotaxis without anesthesia, we identified leaving behavior as a novel behavioral mode in *C. elegans* that emerges after approaching an attractant. The internal state changes driving the transition from approaching to leaving remain unknown, and future studies using genes involved in adaptation, learning, and memory beyond *egl-4* will be essential to uncover its underlying mechanisms.

## Methods

### Caenorhabditis elegans maintenance and strains

*Caenorhabditis elegans* (Bristol N2 strain) were grown under standard conditions (Brenner, 1974). The *egl-4* loss-of-function (MT1073) and *arr-1* loss-of-function (RB660) mutant strains were obtained from the Caenorhabditis Genetics Center.

### Worm preparation for chemotaxis assay

S Basal (1.5 mL) was added to a 6-cm plate, and nematodes were collected into the liquid and transferred to a 1.5-mL tube. After standing for ∼1 min, when adult nematodes had settled to the bottom, the supernatant was removed in step 1. In step 2, S Basal (1.5 mL) was added again, and the sample was pipetted several times. Steps 1 and 2 were repeated three times; in the third wash, deionized water was used to remove S Basal. Approximately 10 µL of liquid containing nematodes was left in the tube, and the remaining supernatant was removed.

### Conventional population chemotaxis assay

Isoamyl alcohol (2 µL; 1:100 dilution) was placed 20 mm from the center of the plate. Ethanol (2 µL) was placed on the opposite side as a control. Worm suspension (10 µL) was applied to the center of the plate, and the gap between the plate and lid was sealed with Parafilm. In the anesthetic condition, sodium azide (1 µL; 1:100 dilution) was placed at the same positions as isoamyl alcohol and ethanol. Time measurement began when the nematodes started moving, and still images were acquired five times at 10-min intervals.

### Center-drop population chemotaxis assay without sodium azide

In the conventional population assay, the attractant is placed 20 mm from the plate center, and the plate edge may constrain approaching individuals. We therefore developed a central-drop method in which the attractant is placed at the center of the plate and nematodes move toward the center. Isoamyl alcohol (2 µL; 1:100 dilution) was placed at the center of the plate. Worm suspension (10 µL) was applied 20 mm from the odor source, and the plate was sealed with Parafilm. Time measurement began when the nematodes started moving, and still images were acquired five times at 10-min intervals.

### Video acquisition of chemotactic behavior

Isoamyl alcohol (2 µL; 1:100 dilution) was placed at the center of the plate. Worm suspension (10 µL) was applied 20 mm from the odor source, and the plate was sealed with Parafilm. The plate was placed on an LED plate and observed under transmitted bright-field illumination. A 50-min video of chemotactic behavior was recorded.

### Quantification of distance from each individual to the isoamyl alcohol source

Using the multi-point tool in the Fiji software, we obtained the coordinates of the isoamyl alcohol source and the coordinates of each nematode at each time point. The isoamyl alcohol coordinates were denoted (x_center, y_center) and each nematode coordinate (x_worm, y_worm). The Euclidean distance between the two points was calculated and then converted from pixels to millimeters using the pixel-per-millimeter scale obtained in Fiji.

### Manual tracking in the population chemotaxis assay

A 15-min chemotaxis video was used. To facilitate visualization, the video was converted from RGB to grayscale in Fiji. Individual trajectories were tracked across all time points using the Manual Tracking plugin, and the resulting trajectories were merged onto the video.

### Development of Simple Worm Tracker

SWT is a custom software tool designed for quantifying single-nematode trajectories. It tracks moving objects using background subtraction. We implemented the software tool in Python and employed the background subtraction model cv2.createBackgroundSubtractorMOG2() from the OpenCV library. When a change of 21 pixels or more was detected between frames, SWT automatically selected the largest contour, computed its centroid, and output the time-series position coordinates. Tracking accuracy can be optimized by adjusting the minimum pixel change threshold based on video and target sizes. The code is available on GitHub https://github.com/ShioriOnoue/SimpleWormTracker.git.

### Trajectory quantification during chemotaxis

Chemotaxis videos were analyzed with SWT to obtain time-series *x-* and *y-*coordinates. Trajectories were plotted from *x-* and *y-*coordinates, with time progression indicated by a color bar (blue at the start, brown at the end).

### Quantification of distance *d* from the nematode to the isoamyl alcohol source

Using the coordinate data obtained above, we quantified the Euclidean distance of each nematode from the isoamyl alcohol source at each time point. Distances were converted from pixels to millimeters using Fiji Set Scale. Ethanol, which does not elicit attraction or avoidance in nematodes, was used as a solvent control.

### Development and validation of a manual sharp-turn selection tool

To quantify run duration and sharp-turn frequency (i.e., significant changes in trajectory angle), we developed a sharp-turn detection tool (Fig. S5). This tool visualizes the SWT trajectory coordinates and allows the user to click on points where sharp turns occur, recording the corresponding coordinates and time points.

To validate whether the tool correctly identifies sharp turns, we quantified sharp-turn frequency in both control runs and isoamyl alcohol–leaving runs using two approaches: (1) manual scoring of sharp-turn times by visual inspection of raw videos, and (2) analysis of the same videos using SWT and the sharp-turn detection tool (Fig. S6). Sharp-turn frequency did not differ significantly between the two approaches in either condition. Thus, the sharp-turn selection tool was used for selecting sharp turns.

### Defining leaving behavior

Initial visual inspection suggested that runs during leaving behavior were longer than those observed during free-moving behavior. We initially defined leaving behavior as runs in which *C. elegans* individuals approached within 5 mm of the isoamyl alcohol source and subsequently moved continuously to a position at least 15 mm away.

To evaluate the robustness of this definition, we analyzed all runs that began after individuals reached the odor source and moved at least 5 mm away. For each run, we measured the maximum distance from the source (“leaving distance”) and calculated sharp-turn frequency as the inverse of run duration. We then plotted leaving distance against sharp-turn frequency and compared leaving runs with free-moving runs, binning leaving distance in 1-mm increments.

Across conditions, when leaving distance was ≥ 6 mm, the isoamyl alcohol condition exhibited significantly lower sharp-turn frequency than the free-moving condition. In the free-moving condition, the distribution of sharp-turn frequency remained relatively stable as the leaving distance increased. In contrast, in the isoamyl alcohol condition, runs with higher sharp-turn frequencies were progressively excluded as the leaving distance increased beyond 6 mm. At a leaving distance of 7 mm, sharp-turn frequency was significantly lower than in free-moving behavior.

Based on these results, we redefined leaving behavior as runs that originated within 5 mm of the odor source and reached at least 7 mm away, as this criterion provided greater robustness across conditions than the initial 15 mm threshold.

### Definition of run duration, leaving distance, and leaving trajectories

Run duration was defined as the elapsed time from one sharp turn to the next sharp turn. Leaving distance was defined as the Euclidean distance between the position at which the nematode was farthest from the isoamyl alcohol source during a run and the coordinates of the source. Leaving trajectories were defined as runs that originated within a 5-mm radius of the isoamyl alcohol source (origin) and reached a position at least 7 mm from the origin.

### Definition of free-moving behavior

Free-moving trajectories were defined as runs during unstimulated behavior that originated within a 5 mm radius of the plate center (origin) and reached a position at least 7 mm from the origin.

### Definition of high-concentration isoamyl alcohol avoidance behavior

Undiluted isoamyl alcohol (2 µL) was placed at the center of an NGM plate. A starting point 5 mm from the odor source was used. A single nematode was transferred to the starting point using a platinum wire pick, and recording began immediately. A 5-min video was acquired and analyzed with SWT. Using the trajectory data, the segment from the starting point to the next sharp turn was defined as high-concentration isoamyl alcohol avoidance behavior.

### Definition of approaching behavior

Isoamyl alcohol (2 µL; 1:100 dilution) was placed at the center of an NGM plate. A starting point 20 mm from the odor source was used. A single nematode was transferred to the starting point using a platinum wire pick, and recording began immediately. A 10-min video was acquired and analyzed with SWT. Using the trajectory data, the segment from the starting point to the next sharp turn was defined as approaching behavior.

### Calculation for locomotor features

#### Sharp-turn frequency (Figs. 3–5)

Using the extracted trajectories, runs were extracted with the sharp-turn selection tool. After quantifying run duration (s) from start to end, we took the inverse to obtain sharp-turn frequency (events/s).

#### Trajectory length (Figs. 3–5)

Before computing trajectory length, trajectories were smoothed to reduce the influence of sinusoidal body bending on positional coordinates. For data recorded at 23.9 frames per second, we applied a moving average with a window size of 45 frames. For each time point, *x*- and *y*-coordinates were replaced by the average across ±45 frames to reduce coordinate jitter. Trajectory length was calculated as the sum of Euclidean distances across all frames.

#### Average speed (Figs. 3–5)

After the same preprocessing as for trajectory length, Euclidean distance per frame and instantaneous speed were calculated, and average speed was then calculated across all frames.

### Statistical analysis

Normality was assessed using the Shapiro–Wilk test (α = 0.05). Homogeneity of variance was evaluated using Bartlett’s test. If data were normally distributed and variances were equal, Student’s *t*-test was used to compare group means. If equal variance could not be assumed, Welch’s *t*-test was used. If data were not normally distributed, the Mann–Whitney *U*-test was used.

### Quantification of the frequency of leaving behavior

For wild-type (N2), *egl-4* loss-of-function (MT1073), and arr-1 loss-of-function (RB660) individuals, 50-min chemotaxis trajectories toward isoamyl alcohol were obtained. The Euclidean distance from each nematode to the isoamyl alcohol source was quantified across all time points. Individuals were classified as “approaching” if they approached the odor source, “leaving” if they exhibited leaving behavior at least once after approaching, or “staying” if they remained near the source. The composition of these categories was displayed as stacked bar graphs. Leaving behavior was defined as a run that continuously moved from within 5 mm of the odor source to a position at least 7 mm away, as defined in N2 individuals.

## Data availability

The datasets are available at ssbd-repos-000504, https://doi.org/10.24631/ssbd.repos.2026.04.504 in SSBD:repository (Kyoda et al., 2025).

## Acknowledgments

We thank Dr. Chentao Wen, Dr. Soya Shinkai, Dr. Ko Sugawara, Dr. Hiroya Itoga, Dr. Go Shioi, Dr. Akiko Hatakeyama, and Dr. Yuki Yamagata for their discussions. We also thank Hatsumi Okada, Rie Furushima, and Tomoko Sugimoto for their kind technical support. An AI-based language editing tool (ChatGPT, OpenAI, Grammarly) was used to improve the clarity and readability of the manuscript. The *Caenorhabditis elegans* strains were provided by the Caenorhabditis Genetics Center, which is funded by the National Institutes of Health (NIH) Office of Research Infrastructure Programs (P40 OD010440).

## Contributions

Onoue S. performed the experiment and computational analysis. Onoue S. and Kyoda K. produced the tracking software. Onoue S. and Onami S. designed the research. Onoue S. and Onami S. wrote the paper. All authors read and approved the final manuscript.

