## Supplemental_information for "Immobilization-free chemotaxis analysis reveals the novel behavioral mode of “leaving” in *Caenorhabditis elegans*"

Supplementary Video 1. This video is available in SSBD:repository at https://doi.org/10.24631/ssbd.repos.2026.04.504. This video corresponds to Figure 2B and shows the central-drop chemotaxis assay of a single *C. elegans*. The video shows the first 5 min of the 50-min recording. The animal starts from the black dot on the right side of the plate.

Figure S1

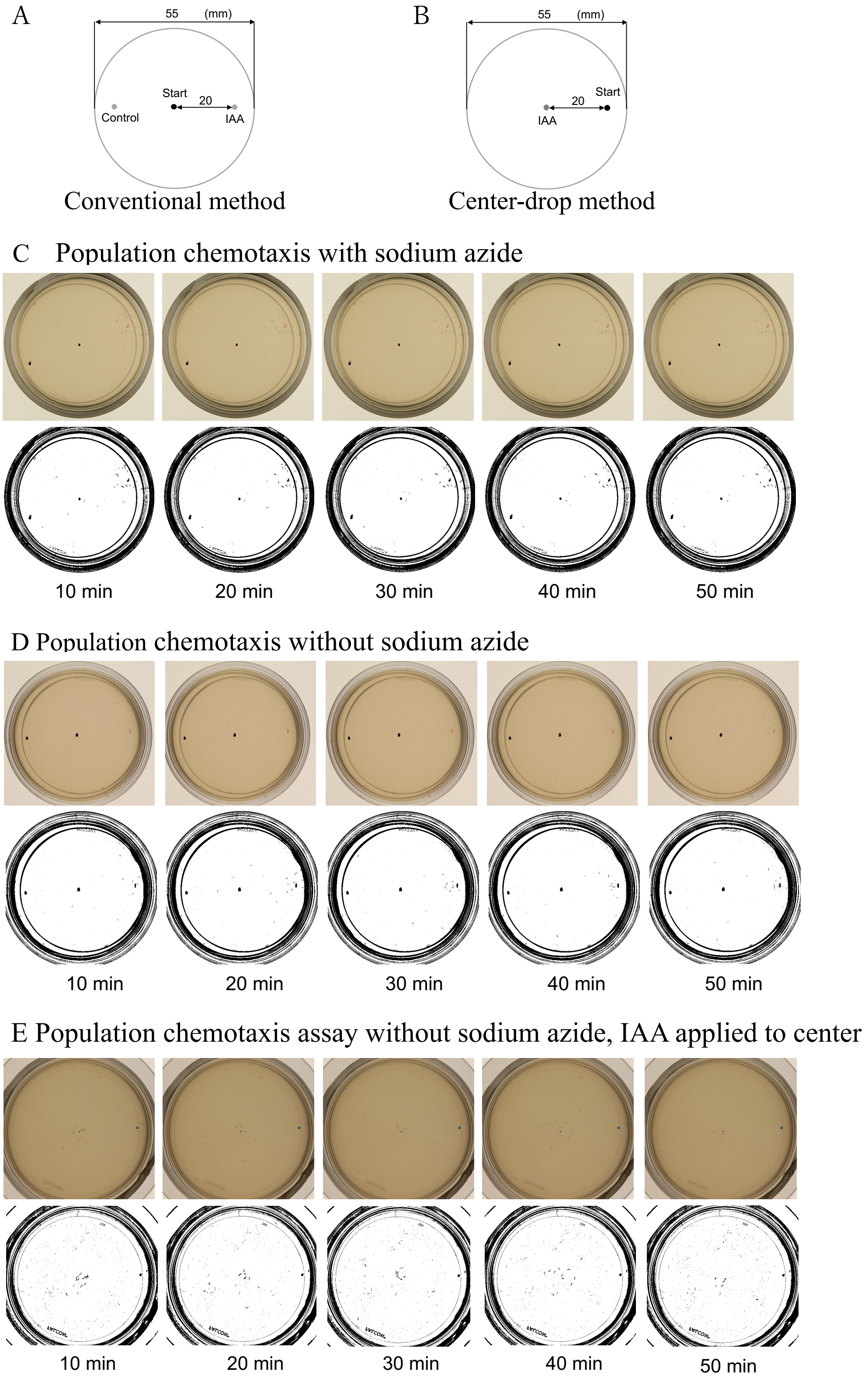
A
55 (mm)
Start
20
Control
IAA
Conventional method
B
55 (mm)
20
Start
IAA
Center-drop method
C Population chemotaxis with sodium azide
10 min
20 min
30 min
40 min
50 min
D Population chemotaxis without sodium azide
10 min
20 min
30 min
40 min
50 min
E Population chemotaxis assay without sodium azide, IAA applied to center
10 min
20 min
30 min
40 min
50 min

Figure. S1. A modified chemotaxis assay enables continuous observation of nematode behavior after reaching an attractant. (A) Conventional chemotaxis assay method. (B) Center-drop chemotaxis assay method: An attractant is dropped in the center of the plate without sodium azide. (C) Conventional chemotaxis assay (C) with sodium azide and (D) without sodium azide. (E) Center-drop method chemotaxis assay without sodium azide. The upper panels displays raw images, while the lower panels were created to facilitate visualization of the nematode's position; the images were binarized and contrast-enhanced.

Figure S2

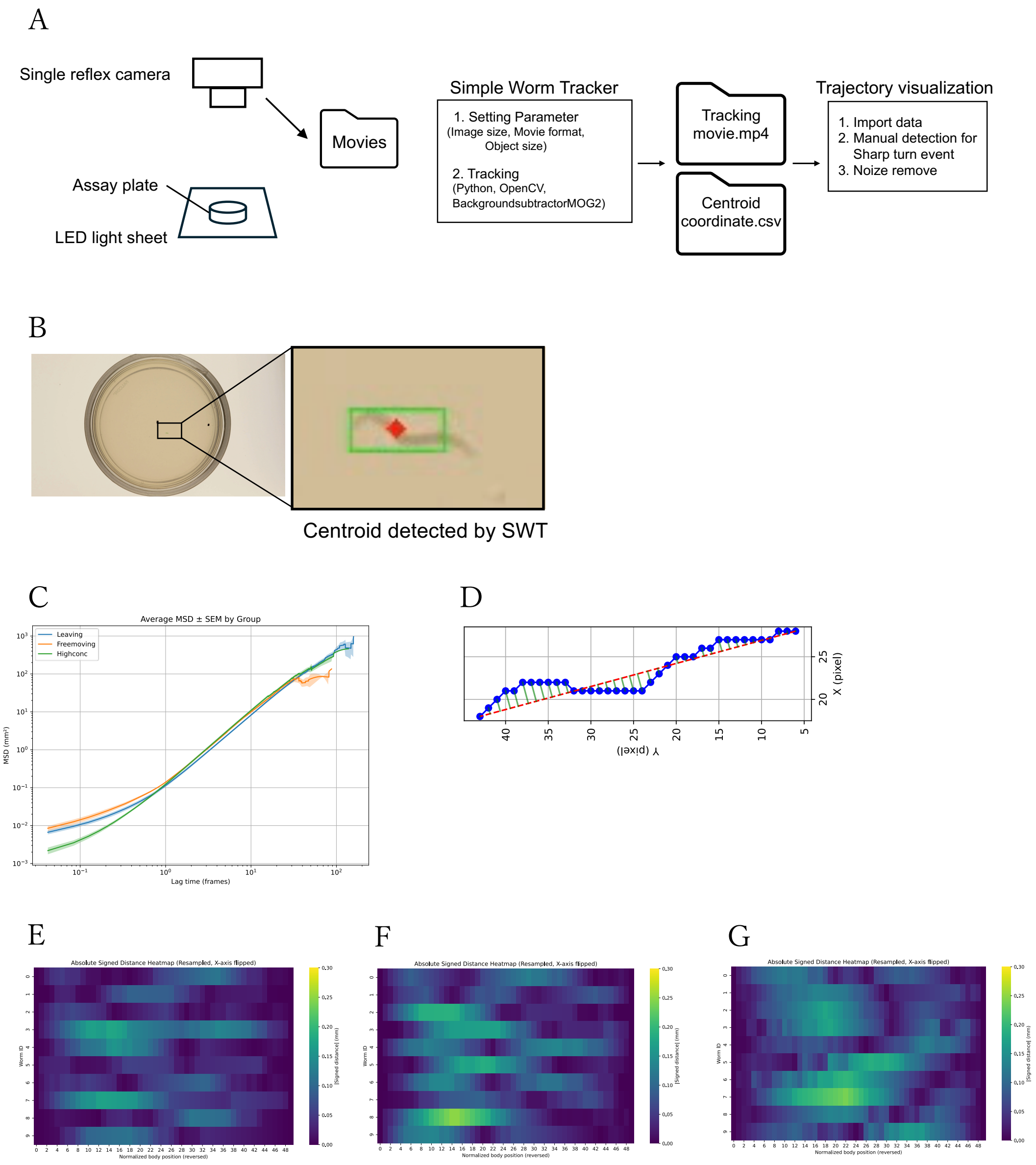


Figure S2. Schematic of the nematode behavior recording and quantitation system. (B) Behavior quantification system used to detect the centroid of the nematode. (C) Mean square displacement (MSD) result for each behavior. (D) Individual nematode's body bending calculation. Body bending result for (E) leaving behavior, (F) free movement, and (G) high-concentration isoamyl alcohol avoidance.

Figure S3

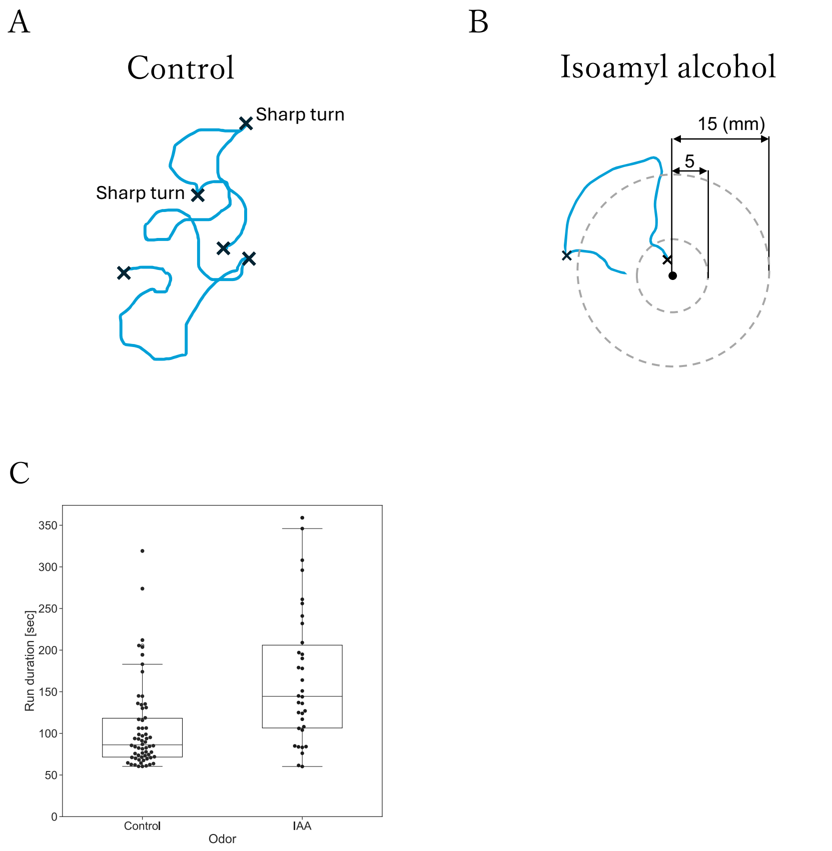


Figure S3. Comparison between leaving and free movement. (A) The run extraction method for free moving as a control. Black crosses indicate sharp turns, and blue line indicates the run trajectory. (B) Method for extracting isoamyl alcohol (IAA) chemotaxis leaving trajectory (i.e., run started from less than 5 mm from the IAA point (black dot) and continuously moving to point more than 15 mm away). (C) Swarm plot of run duration for control (free-moving run) and the IAA leaving trajectory.

Figure S4

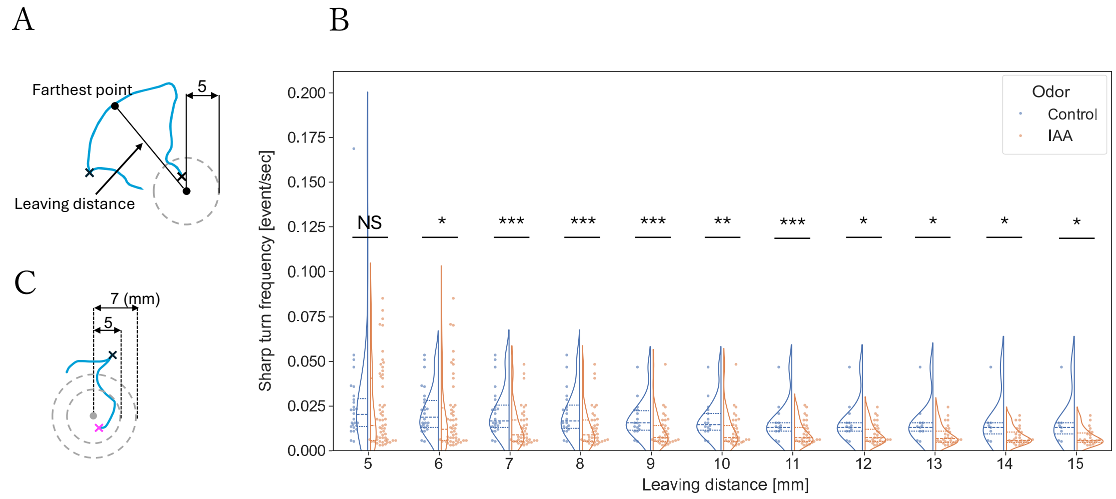


Figure S4. Criteria and analysis flow for defining leaving behavior. (A) Leaving distance: The distance of the nematode from the isoamyl alcohol drop point was defined as the linear distance (mm) between the coordinates of the isoamyl alcohol drop point and the coordinates of the time when each nematode was farthest from the isoamyl alcohol drop point within a run. (B) Each run’s leaving distance and sharp-turn frequency were compared with free-movement runs and leaving. Free movement is shown in blue, leaving in orange. Significance of the Mann-Whitney U-test results for the sharp-turn frequency for each group are noted as **p* < 0.05, ***p* < 0.01, ****p* < 0.001. (C) Leaving behavior: Runs with a starting point within a 5-mm radius of the origin at the isoamyl alcohol drop point, reaching a radius of at least 7 mm from the origin.

Figure S5

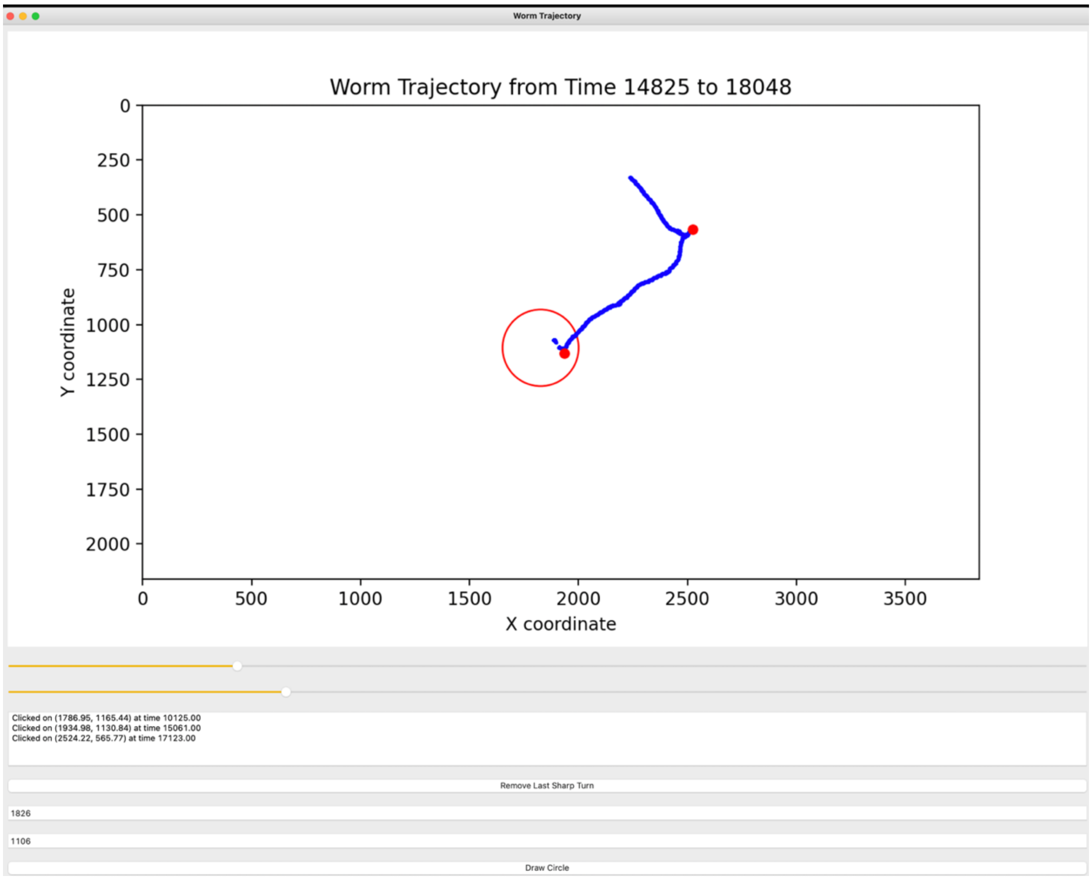


Figure S5. We developed a sharp-turn selection tool to quantify the run duration and frequency of sharp turns. This system visualizes the action trajectory coordinate data quantified by Simple Worm Tracker. By visually selecting and clicking on the point where a sharp turn occurs and a dramatic angle change occurs, the coordinates and time point of the clicked trajectory data can be obtained. The system also incorporates a graphic drawing module that displays a circle with a radius of 5 mm in red, centered at the point where the isoamyl alcohol drop occurs. By selecting a run that falls outside this circle, the leaving behavior can be efficiently extracted.

Figure S6

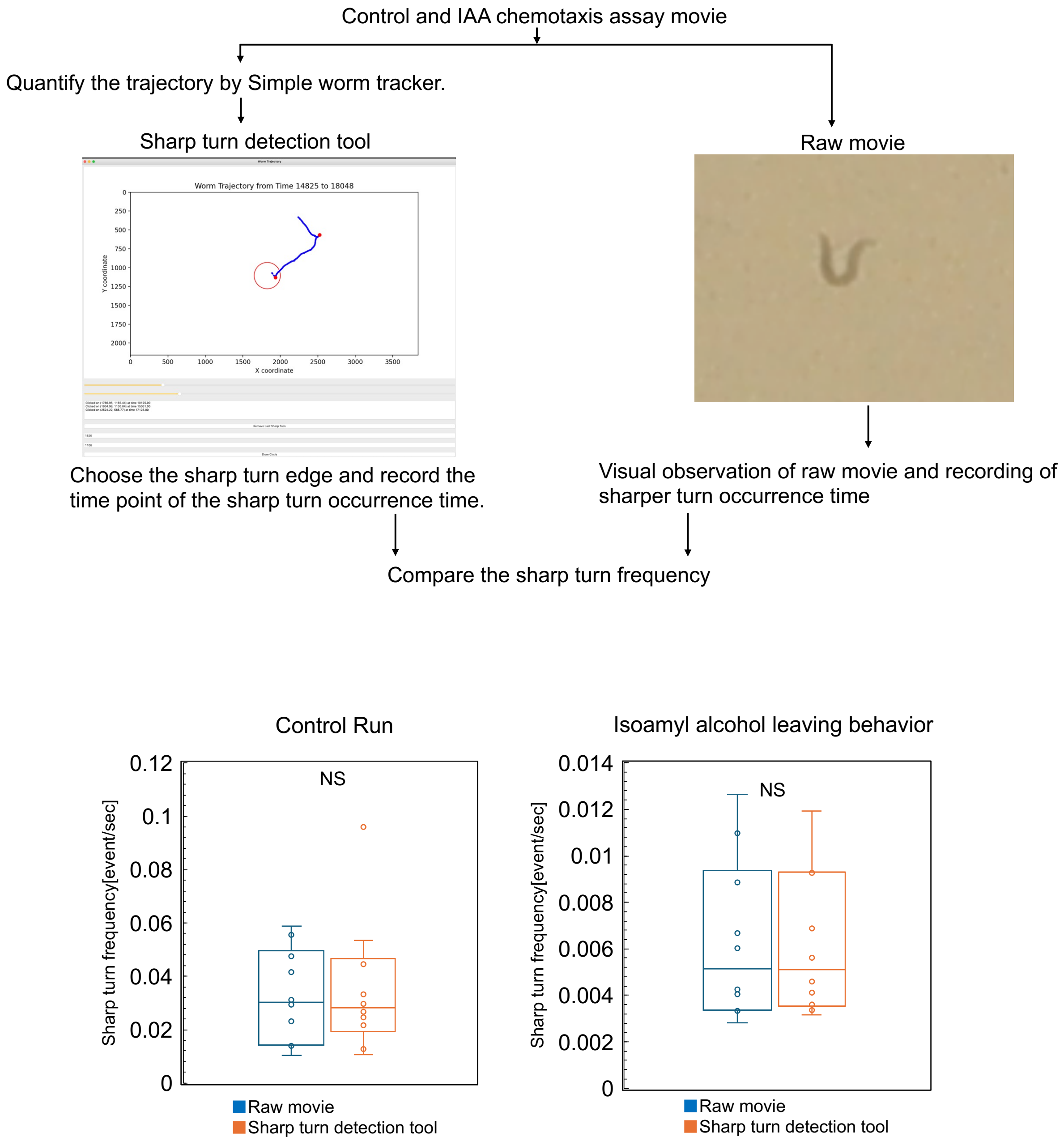


Figure S6. Performance evaluation of the sharp-turn detection tool. To confirm whether the tool correctly selected the sharp-turn events, we compared two methods: (1) visually observing raw movies of both control run and leaving and recording the time of sharp-turn occurrence and (2) obtaining the time of sharp-turn occurrence by analyzing the same movies with Simple Worm Tracker and the sharp-turn detection tool. The sharp-turn frequency was quantified and compared with that obtained by analyzing the same videos using the Simple Worm Tracker and the sharp-turn detection tool. The results showed no significant difference in sharp-turn frequency between visual observation of the raw movie and using the sharp-turn selection tool for both conditions, indicating the sharp-turn selection tool is effective. Control: raw movie (n = 7), sharp-turn detection tool (n = 8); isoamyl alcohol: raw movie (n = 7), sharp-turn detection tool (n = 8). Correspondence to Shuichi Onami.
